# A burst of transposon expression accompanies the activation of Y chromosome fertility genes during Drosophila spermatogenesis

**DOI:** 10.1101/2021.05.10.443472

**Authors:** Matthew A. Lawlor, Weihuan Cao, Christopher E. Ellison

**Affiliations:** Department of Genetics, Human Genetics Institute of New Jersey, Rutgers University, Piscataway, New Jersey

## Abstract

Transposable elements (TEs) must replicate in germline cells to pass novel insertions to offspring. In *Drosophila melanogaster* ovaries, TEs can exploit specific developmental windows of opportunity to evade host silencing and increase their copy numbers. However, TE activity and host silencing in the distinct cell types of the *Drosophila melanogaster* testis are not well understood. We reanalyzed publicly available single-cell RNA-seq datasets to quantify TE expression in the distinct cell types of the *Drosophila* testis. We developed a novel method for identification of TE and host gene expression programs and find that a distinct population of early spermatocytes expresses a large number of TEs at much higher levels than other germline and somatic components of the testes. This burst of TE expression coincides with the activation of Y chromosome fertility factors and spermatocyte-specific transcriptional regulators, as well as downregulation of many components of the piRNA pathway. The TEs expressed by this cell population are enriched on the Y chromosome and depleted on the X chromosome relative to other active TEs. These data suggest that some TEs may achieve high insertional activity in males by exploiting a window of opportunity for mobilization created by the activation of spermatocyte-specific and Y-chromosome-specific transcriptional programs.

## Introduction

Transposable elements (TEs) are abundant in the genomes of plants and animals despite the presence of sophisticated host genome defense pathways. The genetic mechanisms responsible for the evolutionary success and persistence of TEs remain unclear. It is possible that the fitness benefit of complete TE suppression is not large enough to be evolutionarily favorable (Charlesworth and Langley 1986; Lee and Langley 2010; Kelleher and Barbash 2013). On the other hand, it is also possible that, like many viruses, TEs are engaged in an evolutionary arms race with their hosts, with TEs continuously evolving to escape silencing and the host genome continuously evolving to reestablish TE suppression (Parhad and Theurkauf 2019). Many host genes involved in TE defense are rapidly evolving, consistent with ongoing host-TE conflict (Kolaczkowski, Hupalo, and Kern 2011; Obbard et al. 2011, 2006; Simkin et al. 2013; Helleu and Levine 2018; Crysnanto and Obbard 2019), however relatively few strategies where TEs can escape or evade host silencing have been identified (Cosby, Chang, and Feschotte 2019). In the Drosophila ovary, there is evidence that some TEs propagate in permissive nurse cells and hijack the host’s mRNA transport pathway to move to the developing oocyte, which is more recalcitrant to TE expression (Wang et al. 2018). In another study, Dufourt et al. identified a small region of mitotically dividing germline cysts where the piRNA pathway gene Piwi is depleted and TE silencing is much weaker than in the surrounding cells. They termed this region the “piwiless pocket” and proposed that TEs may take advantage of this niche to replicate in the Drosophila germline (Dufourt et al. 2014).

TE replication and host silencing have been extensively studied in the Drosophila ovary, however surprisingly little is known about these same phenomena in the testes. Several previous observations suggest that there may be substantial differences between ovaries and testes with respect to both TE activity levels and host silencing pathways. For example, multiple TE families are known to exhibit strong sex biases: The *I-element, P-element*, and *gypsy* TE families are all expressed at higher levels in the female germline (Busseau et al. 1994; Pélisson et al. 1994; Roche, Schiff, and Rio 1995) whereas the opposite is true for the *copia, micropia, 1731*, and *412* TE families (Lankenau, Corces, and Lankenau 1994; Haoudi et al. 1997; Pasyukova et al. 1997; Borie et al. 2002). The piRNA pathway is active in both somatic and germline cells in the ovary and piRNAs bound by *Aub* and *Ago3* undergo robust ping-pong amplification in the ovarian germline. In the testes, TE-derived piRNAs are produced in germ cells, however the vast majority (~75%) arise from the *suppressor of stellate* [*Su(Ste)*] and *AT-chX* satellite repeats, rather than the canonical piRNA clusters that have been identified in ovaries (Quénerch’du, Anand, and Toshie 2016; P. Chen et al. 2020). Furthermore, many TE families show large differences in piRNA abundance between ovaries and testes (P. Chen et al. 2020) and TE-derived piRNAs only show a weak signature of ping-pong amplification in spermatocytes, likely due to low levels or absence of *Ago3* (Quénerch’du, Anand, and Toshie 2016).

Here we have analyzed TE expression at single-cell resolution in order to gain insight into the dynamics of TE activity in Drosophila testes. We develop a novel approach for identification of TE and host gene expression programs and find that a subset of primary spermatocytes expresses a diverse group of TEs at high levels relative to other cell types. These TEs are co-expressed with Y-linked fertility factors and we find evidence that they are more active in males compared to females. These data suggest some TEs may exploit spermatocyte-specific transcriptional programs and Y chromosome activation to remain active in the *Drosophila melanogaster* genome.

## Results

### Data processing and cell type identification

We reanalyzed 10x Genomics 3’ single-cell expression data from a recent study examining sex chromosome gene expression in *D. melanogaster* larval testes (Mahadevaraju et al. 2020). The Drosophila larval testes are elongated spheres encased in epithelial cells. Their apical caps contain germline stem cells and the somatic cells of the GSC niche, the hub cells. The apical caps of the testes house mitotically dividing spermatogonial cysts, while the middle portion houses meiotic spermatocyte cysts encased by pairs of somatic cyst cells. L3 larval testes include germ cell stages from GSC through primary spermatocytes, which exist in an extended meiotic prophase.

To quantify transposable element expression at single-cell resolution, we masked TE sequences in the *Iso1* D. melanogaster release 6 genome assembly and appended the consensus sequences for all *D. melanogaster* RepBase TEs (Bao, Kojima, and Kohany 2015). We used this custom reference sequence to generate an aligner index for the 10x Genomics Cell Ranger 3.1.0 single-cell expression alignment and quantification pipeline (Zheng et al. 2017). We used scrublet (Wolock, Lopez, and Klein 2019) to remove putative doublet barcodes and applied scanpy (Wolf, Angerer, and Theis 2018) for basic preprocessing, normalization, scaling, and merging of the replicate datasets. To identify transcriptionally similar cell clusters, we excluded all transposons and generated a nearest neighbors graph. We applied the Leiden algorithm (Traag, Waltman, and van Eck 2019) to reveal 10 clusters, including several highly distinct clusters and several clusters with high degrees of similarity, as indicated by adjacency in the UMAP embedding (Figure 1A). We classified each of these clusters using garnett (Pliner, Shendure, and Trapnell 2019) and a collection of curated markers (see Methods, Supplementary Table 1) to assign each cell to a known testis cell type (Figure 1B).

**Figure 1:**
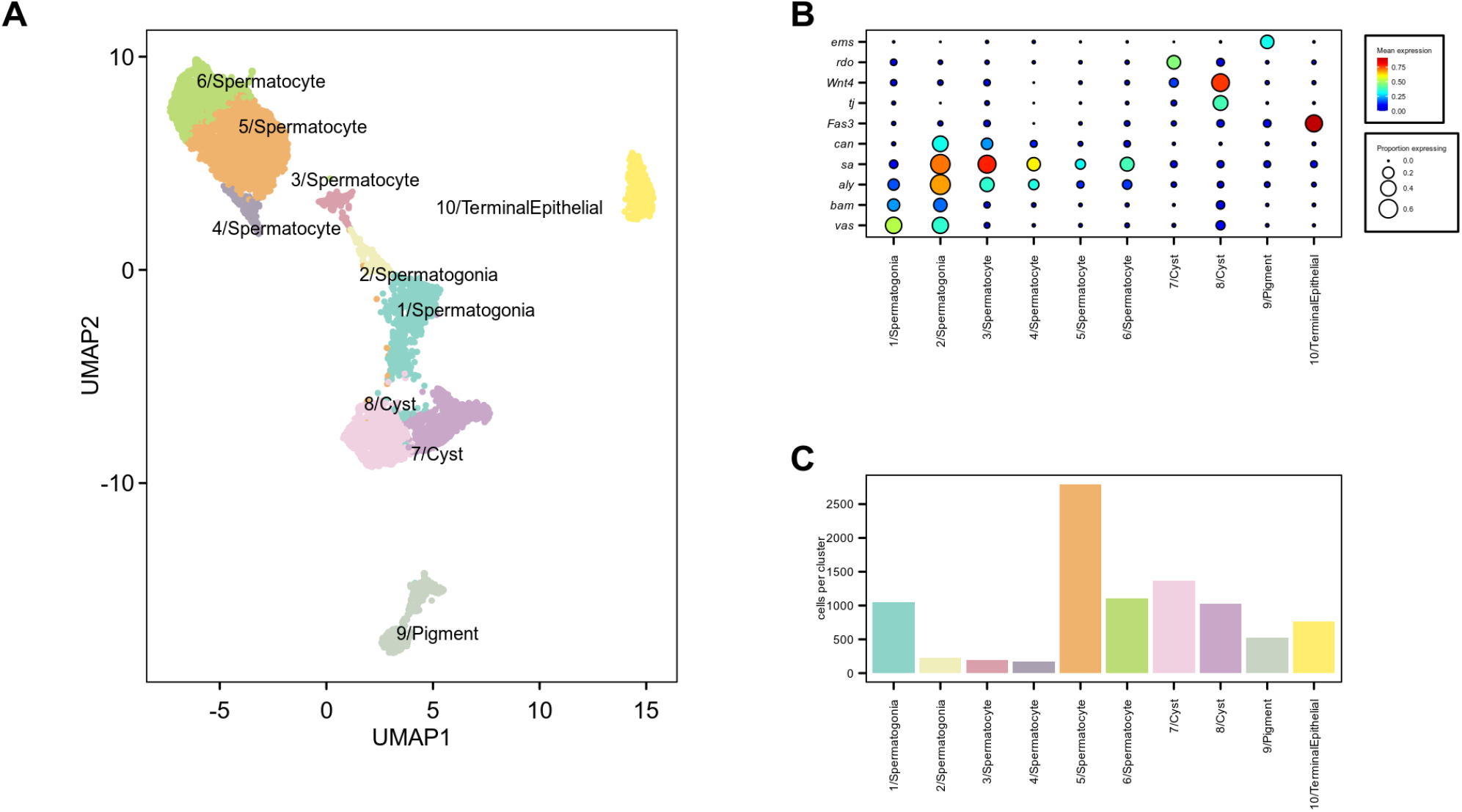
Identification of testis cell populations. **A)** UMAP projection groups transcriptionally similar cells in 2D space. Cells are colored by assigned cell type. **B)** Dot plot shows expression of selected marker genes used for cell type assignment. Color of each dot corresponds to mean normalized and log-transformed expression within cell clusters. Dot size corresponds to the proportion of cells in each cluster expressing the marker. **C)** Cell counts within each cell type cluster.

Our filtering approach is more conservative than applied to these data in their initial study, yielding a final dataset with fewer cells than originally published (Supplementary Figure 1D). To assess the similarity of our clusters with previously published clusters, we generated mean expression values per cluster for each gene and computed Spearman’s rank correlation for each pairwise combination of clusters from each study (Supplementary Figure 1C). There is a strong correspondence between our clusters and those previously identified from these data, with high pairwise correlations for every cluster previously reported, though minor differences are apparent. Notably, we identify fewer distinct cyst cell clusters but more distinct spermatocyte clusters than reported in the original study.

Similar to the previously described analysis of these data (Mahadevaraju et al. 2020), we identify distinct somatic and germline clusters (Figure 1A). We identify cyst cells (clusters 7 and 8) which express *tj* and *wnt4* at high levels. Hub and terminal epithelial cells (cluster 10) are defined largely by *Fas3* expression, and pigment cells (cluster 9) express *Sox100B* (Figure 1B).

The remaining cells comprise the germline components of these data. Cluster 1 contains germline stem cells and early spermatogonia, marked by *vasa* and *spn-E* (Figure 1B). A second spermatogonial cluster (2/Spermatogonia) expresses spermatogonial markers such as *bam* and spermatocyte markers such as *aly*, which respectively are required for GSC differentiation and initiation of a primary spermatocyte transcription program. They are most transcriptionally similar to the G spermatogonia cluster identified by Mahadevaraju *et al.* but mean normalized UMI counts for this cluster also correlate well with that study’s E1 early spermatocyte cluster (Supplementary Figure 1C). This observation suggests that our cluster 2 may represent spermatogonia just beginning the transition to meiotic prophase or the very early spermatocytes.

The final four clusters (3, 4, 5, and 6) represent the majority of filtered cells (Figure 1C) and express *aly* as well as *sa* and *can*, which are effectors of the primary spermatocyte expression program (Beall et al. 2007; White-Cooper et al. 1998) (Figure 1B). These clusters are transcriptionally similar to primary spermatocytes identified previously (Supplementary Figure 1C). Mean expression in clusters 3 and 4 correlates well with the previously reported early primary spermatocytes while expression in 5 and 6 correlates most highly with previously reported middle and late primary spermatocyte clusters (Supplementary Figure 1C). Taken together, these observations suggest that the germline clusters may be ordered from earliest to latest differentiation state by the cluster numbers reported here. However, among the later putative spermatocyte clusters (4, 5, and 6) it is challenging to definitively identify the differentiation order.

### A spermatocyte subpopulation shows high expression of transposable elements

To quantify cell-type-specific TE expression in the *Drosophila* testis, we began by visualizing expression of all TEs with at least 3 UMIs detected across all individual cells (Figure 2). While individual somatic cyst cells and pigment cells sporadically express a small number of TEs, there is no evidence for cell-type specific upregulation of distinct TE families in these cells. On the other hand, a small number of TE families show high expression specifically in the terminal epithelial or spermatogonia clusters. Most striking, however, are the cells from cluster 3 spermatocytes, whose members uniformly express a relatively large number of TE families at high levels (Figure 2). In fact, cluster 3 spermatocytes have the most TE-derived UMIs per cell, for both depth-normalized and raw UMI counts (Supplementary Figure 2A, 2B). UMAP embedding and Leiden clustering was performed on highly variable host genes only, suggesting that cluster 3 is transcriptionally distinct from other spermatocytes independent of TE expression.

**Figure 2:**
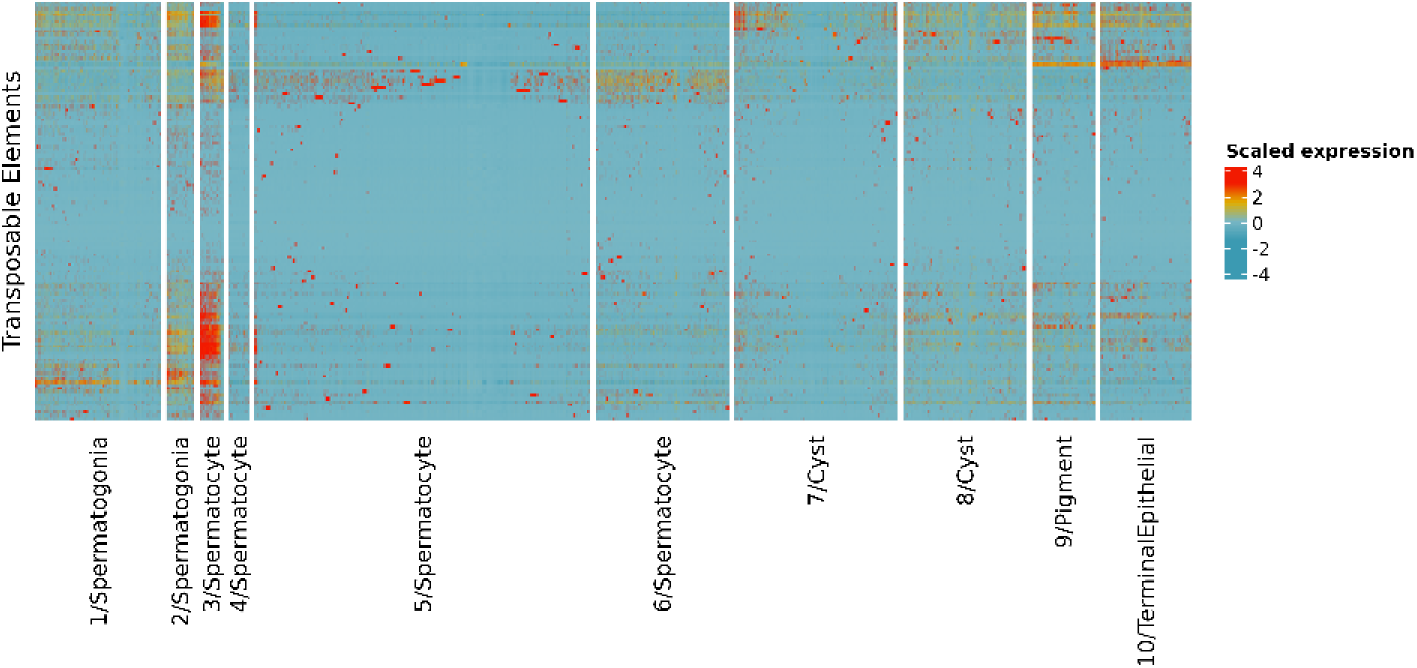
A spermatocyte cluster expresses transposons at high levels. Heatmap shows scaled expression level of all transposable elements detected in this dataset across all cells. Several clusters express small groups of transposable elements uniformly or show sporadic expression of transposons in some member cells. Cluster 3/Spermatocyte shows uniformly high expression of many transposable elements.

To verify that the detected TE expression pattern is not a technical artifact of 10X scRNA-seq, we aligned L3 larval testis poly-A selected RNA-seq reads generated alongside the single cell data (Mahadevaraju et al. 2020). Summarized pseudo-bulk expression estimates derived from the scRNA-seq data are highly concordant with bulk expression both globally and with respect to TEs specifically (Supplementary Figure 1A, 1B). We next assessed whether TE fragments nested in other cellular RNAs may be artificially increasing measurements of TE expression in the testes. While several families (2 out of 125 families analyzed) exhibit extreme coverage at localized portions of their consensus sequence, consistent with truncated copies and/or host gene-TE fusions, the vast majority of TE families expressed in testis show coverage throughout their consensus sequences and within-TE RNA-seq signal variability is comparable to single isoform host genes (Supplementary Figure 2C, 2D). We additionally queried poly-A RNA-seq reads from w1118 testis to test if detected TE expression is a consequence of chimeric transcripts produced by TE insertions within host genes. Only a small number of TEs show evidence of reproducible chimeric transcripts (Supplementary Figure 2E, 2F).

### Independent Component Analysis reveals a TE-enriched gene expression program

To identify host gene expression programs (GEPs) co-expressed with TEs, we implemented a GEP detection pipeline using Independent Component Analysis (ICA). ICA has previously been shown to perform favorably compared to other GEP detection methods (Saelens, Cannoodt, and Saeys 2018). Along with other matrix-factorization approaches, ICA yields a biologically interpretable pair of matrices.

Some factorization approaches, such as ICA and non-negative matrix factorization (NMF) suffer from stochastically varying solutions. Kotliar et al. have previously introduced an elegant approach, termed consensus NMF (cNMF), to stabilize NMF solutions for scRNA-seq GEP detection (Kotliar et al. 2019). This approach clusters the results of many iterations of NMF to buffer the influence of outlier solutions yielded by single runs of the algorithm. However, when we implemented this approach, we found that it yielded large GEPs utilized by broad cell types. We therefore chose to use ICA factorization because this approach was able to group genes into smaller GEPs expressed specifically by smaller cell populations.

We applied this consensus approach to ICA to address the issue of ICA solution randomness. We additionally applied a grid search approach to choose the two most important parameters of our pipeline – the number of components (*k*) to decompose the single cell expression matrix into and the appropriate cutoff (*q*) for identifying the distinct members of each GEP (see Methods).

We ran our consensus ICA approach three times for combinations of selected k-values from 10 to 150 and q-value cutoffs from 0.001 to 0.16. We assessed the biological interpretability of the candidate solutions by enrichment for Gene Ontology Biological Process (GO:BP) terms. We found that optimizing only for maximum percentage of GO:BP-enriched GEPs yielded mostly large GEPs associated with very general biological processes. Under the assumption that a maximally interpretable set of GEPs should capture a wide range of biological processes and should favor discovery of minimally redundant GEPs, we then calculated two scores: one based on the breadth of GO:BP enrichments in a given ICA solution and the other on the unique assignments of GO terms to GEPs (see Methods). These scores are highly reproducible across replicate runs of cICA (Supplementary Figure 2A).

We combined these two metrics into a joint score for each combination of k- and q-parameters. We selected an optimal combination of k (90) and q (0.005) and used the GEPs identified from independent runs as our working GEPs for this study. The GEPs identified using this approach range in size from 10 genes to over 600, with 75% of identified GEPs containing 200 or fewer genes (Supplementary Figure 2B). Sixty-nine percent of identified GEPs were enriched at p < 0.05 for a Biological Process GO term not enriched in any other GEP (Supplementary Figure 3C).

Our method identified many GEPs with at least one TE included alongside host genes, but a single GEP (GEP-27) included over 70 transposons along with approximately 300 host genes (Figure 3A, 3B). All major classes of TEs are represented in this GEP, including LTR and non-LTR retroelements and DNA transposons. Interestingly, we find that these TEs are enriched for elements located within the flamenco piRNA cluster, which is involved in TE suppression in ovarian follicle cells (Brennecke et al. 2007) (Fisher’s Exact Test P=0.03). Several other TEs in this GEP have previously been shown to be male-biased: the LTR retrotransposons *1731, 412*, and *copia* are expressed at high levels in the primary spermatocytes of *D. melanogaster* (Haoudi et al. 1997; Borie et al. 2002; Pasyukova et al. 1997), while *micropia* transcripts have been shown to be associated with Y chromosome lampbrush loops in the primary spermatocytes of *D. hydei* (Lankenau, Corces, and Lankenau 1994). We visualized per-cell expression scores (see Methods) for GEP-27 on the UMAP projection and observed that it is expressed exclusively by cells in cluster 3, in agreement with our visual inspection of TE expression across the dataset (Figure 3C). These results suggest that a burst of TE expression occurs in a distinct subcluster of primary spermatocytes in the larval testes.

**Figure 3:**
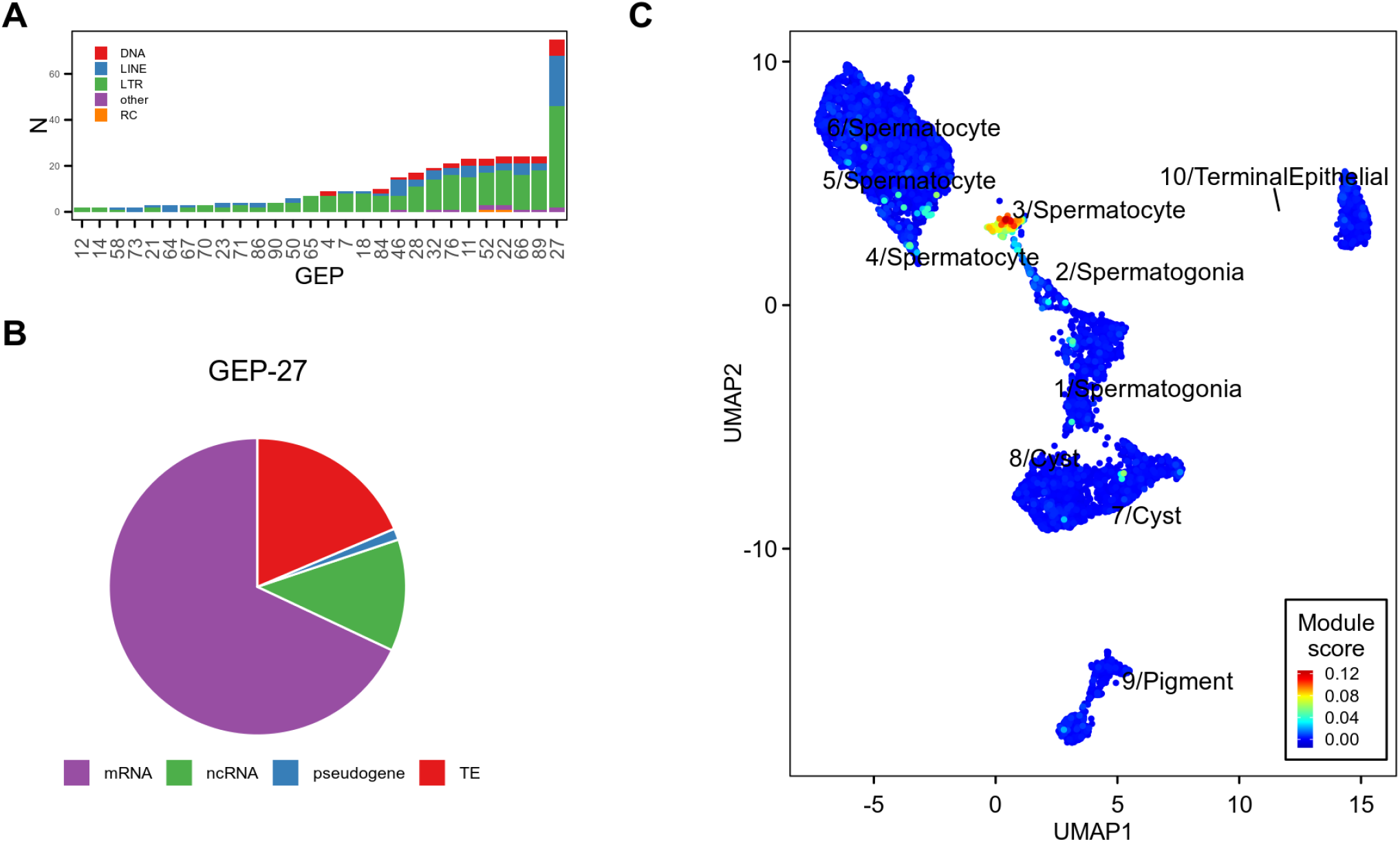
A TE-enriched gene expression program is expressed primarily in cluster 3/Spermatocytes. **A)** Tallies of TE classes found in each GEP containing at least 1 TE. GEP-27 contains almost four-fold more TEs than the next most TE-rich GEP and is predominately composed of LTR retrotransposons (59%%), LINE (29%) and DNA (9%) elements. **B)** GEP-27 contains over 300 total features, the vast majority of which are either protein-coding genes (68%), TEs (18.6%), or non-coding RNAs (12%). **C)** UMAP projection colored by GEP-27 expression score. Expression score is derived from the consensus (averaged) ICA source matrix.

We identified *EAChm*, a host gene TEP member highly expressed in the TEP-expressing population (Figure 4A) as a marker for TEP-expressing spermatocytes. EAChm is an enhancer of *chm* acetyltransferase activity that shows high expression in testis in modEncode RNA-seq data (J. B. Brown et al. 2014; Larkin et al. 2021). Its role in spermatogenesis is currently unknown. We next performed multiplexed RNA-FISH in whole mount L3 testes for *EAChm* and two TEP-TEs, *ACCORD2* and *QUASIMODO2* (Figure 4A) to confirm co-expression of TEP-TEs with host genes in 3/Spermatocyte cells. We find that *EAChm, ACCORD2*, and *QUASIMODO2* show similar spatial patterns of expression, consistent with their membership in the same gene expression program (Figure 4C, Supplementary Figures 5, 6, 7). Furthermore, the transcripts of all three elements are confined to the central portion of the larval testis, in agreement with our assessment that TEP-expressing cells are primary spermatocytes.

**Figure 4.**
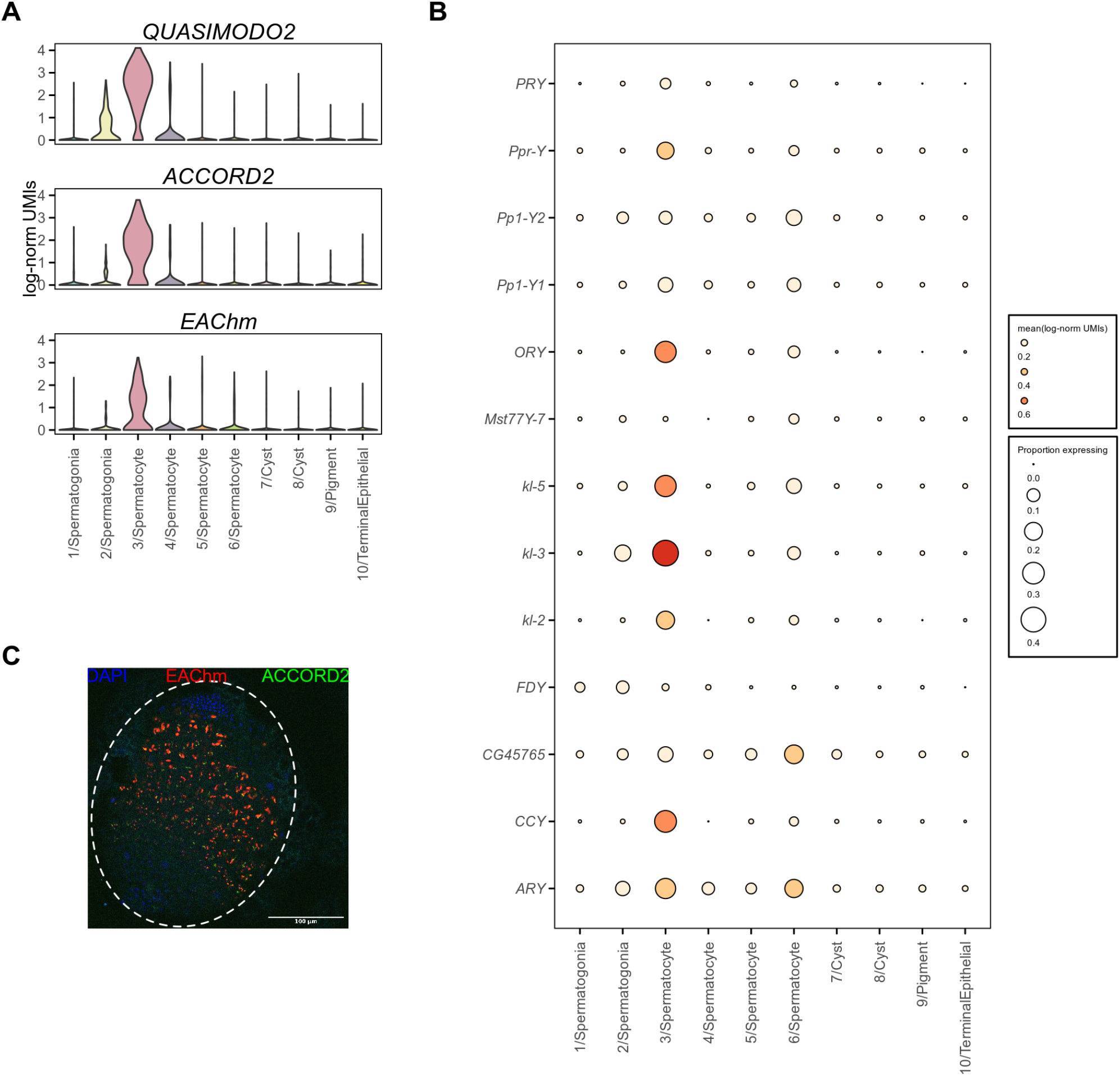
TEP co-occurs with Y chromosome transcriptional activity. **A)** Violin plot shows normalized expression of *EAChm, QUASIMODO2*, and *ACCORD2* is largely confined to cluster 3/Spermatocyte. **B)** Multiplexed RNA-FISH in whole-mount 3rd larval instar w1118 testis shows *ACCORD2* and *EAChm* expression is detected in the middle region of the testis, where primary spermatocytes are located. Red: *EAChm*; green: *ACCORD2*; blue: DAPI. **C)** Dot plot shows mean normalized expression of Y-linked genes in each cluster. Dot size corresponds to the proportion of cells in each cluster with detectable expression of each Y-linked gene. Y-linked genes, especially fertility factors *kl-3* and *kl-5*, are highly expressed by cluster 3/Spermatocyte.

We next sought to determine whether the same gene module is expressed in the testes of adult flies. To do so, we reanalyzed previously published single-cell RNA-seq from adult testes of a different *D. melanogaster* strain (Witt et al. 2019). The TE expression profile from our 3/spermatocyte cells that express GEP-27 is highly correlated with a putative spermatocyte cluster we identified in the Witt et al. data, suggesting that the TEP we identified in larval testes is also expressed in the testes of adults as well as other strains of *D. melanogaster* (Spearman’s R=0.49, P=9.3e-6)(Supplementary Figure 3D).

In order to better understand why TEs are upregulated specifically in the cluster 3 cells, we examined GEP-27 and found that the program contains primary spermatocyte-restricted genes that are required for sperm maturation (Supplementary Figure 8A). Two testis-specific TBP associated factors (TAFs), *can* and *sa*, are members of the TEP, although they are not exclusively expressed in cluster 3. Testis-specific Meiotic Arrest Complex (tMAC) components *aly* and *wuc*, which promote transcription of spermatocyte-specific genes by activating alternative promoters (Lu et al. 2020), are members of the TEP, as well as *kmg*, which blocks promiscuous activation of genes by tMAC (Kim et al. 2017). This supports our analysis suggesting cluster 3 is predominantly composed of primary spermatocytes.

GO enrichment analysis shows that GEP-27 is enriched for genes that function in axonemal assembly and cilium movement, including the Y chromosome fertility factors *kl-2, kl-3*, and *kl-5*, which are expressed specifically in primary spermatocytes (Goldstein, Hardy, and Lindsley 1982)(Figure 4C, Supplementary Figure 8B). GEP-27 is significantly enriched for genes from the Y chromosome: 6 of 9 Y chromosome genes detected in these data are assigned to GEP-27 (Supplementary Figure 8C, Chi-square test P=1.7e-05).

In meiotic prophase, 16-cell primary spermatocytes undergo chromatin decondensation and greatly increase in size (McKee, Yan, and Tsai 2012). Y-chromosome lampbrush loops also form at this stage of development and the Y chromosome becomes enriched for the H3K9ac histone modification, which is associated with active transcription (Hennig and Weyrich 2013). Consistent with this phenomenon, we also find that *tplus3a* and *tplus3b*, two genes required for expression of Y chromosome fertility factors (Hundertmark et al. 2019), are members of GEP-27 as well as *bol*, which binds the decondensed giant introns of several Y loop-forming genes (Redhouse, Mozziconacci, and White 2011). Expression of the 6 Y-linked genes, *bol, tplus3a*, and *tplus3b*, is highest in cluster 3 spermatocytes (Supplementary Figure 8A). These results are consistent with the burst of TE activity that we observe in cluster 3/Spermatocyte cells coinciding with the activation of the Y chromosome fertility genes.

### TEP-TEs are enriched on the Y chromosome

Given that the TEP-TEs are co-expressed with Y chromosome fertility genes, we hypothesized that their upregulation is due to activation of Y-linked copies of these TEs. To address this hypothesis, we first investigated whether TEP-TEs do indeed have copies that are located on the Y chromosome.

We first used RepeatMasker to identify transposon insertions in a recently published *Drosophila melanogaster Iso1* strain genome assembly with improved Y chromosome content (Chang and Larracuente 2019) compared with the current *D. melanogaster* Release 6 reference sequence. We found that 70% of TEP-TEs have at least one full-length copy located on a known Y-linked scaffold (Supplementary Figure 8D) and a significantly larger percentage of TEP-TE insertions are found on the Y chromosome compared with other expressed TEs (Chi-square test P= 2.29e-292, Figure 5A). We also estimated male-specific TE copy numbers by performing Illumina whole-genome sequencing (WGS) of males and females from strain *w1118*. We found that TEP-TEs have significantly elevated copy numbers in males, compared to females, as expected if these TEs have insertions located on the Y chromosome (Wilcoxon Rank-Sum test P=0.0035, Figure 5B).

**Figure 5.**
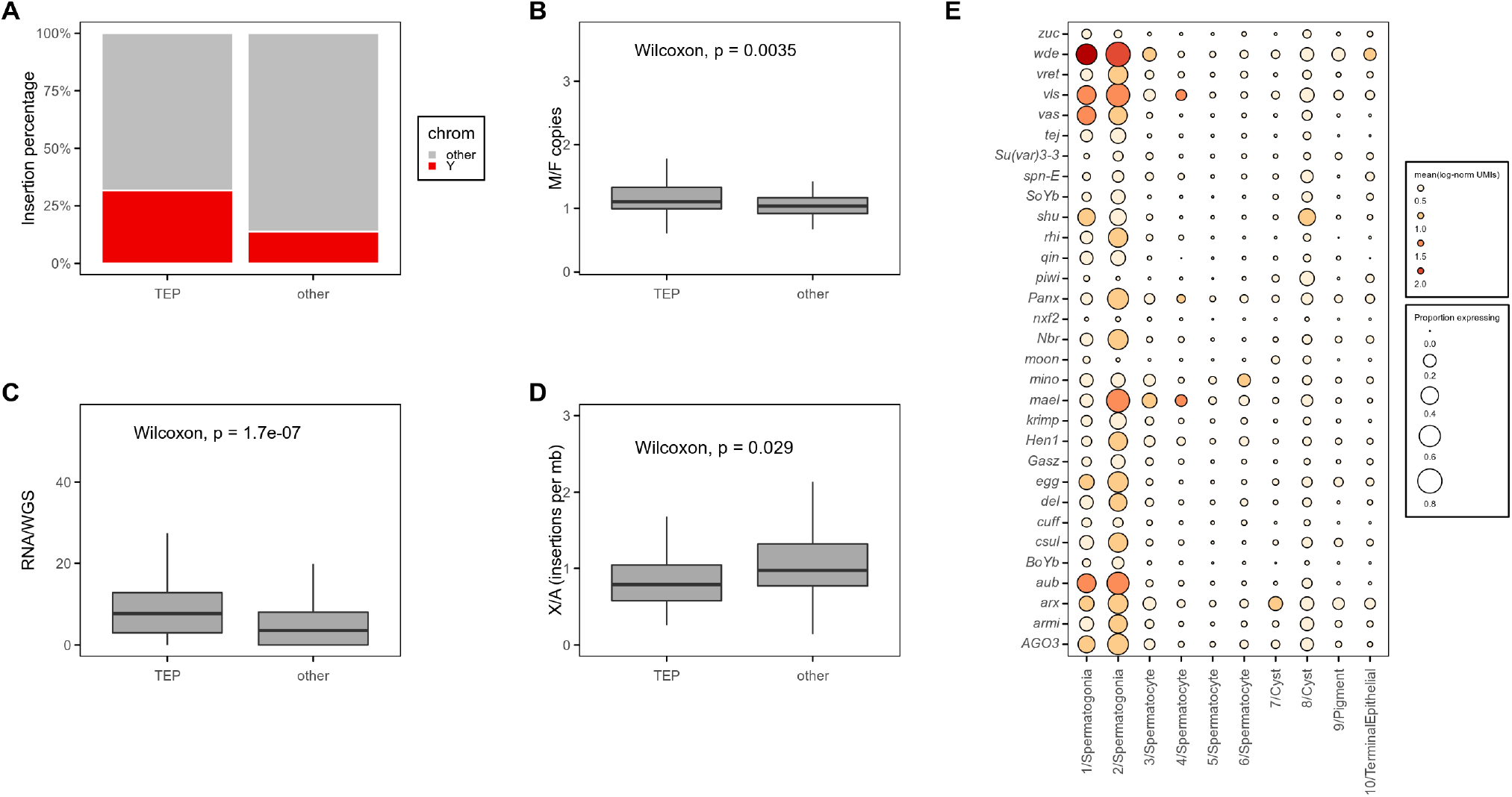
TEP-TEs are enriched on the Y chromosome. **A)** A higher proportion of TEP-TE insertions are found on the Y chromosome compared to non-TEP-TEs. Chi-square test, P = 2.29e-292. To better estimate Y-linked insertions despite the incomplete Y-assembly in the reference sequence, insertions were mapped from a heterochromatin-enriched assembly (see **Methods**). **B)** Ratios of male and female copy number for individual TEs estimated from w1118 WGS coverage. TEP-TEs are present at higher copy number in the male genome Wilcoxon rank-sum test, P=0.0035. **C)** Allele specific analysis of TE expression (see **Methods**) shows that Y-linked copies of TEP TEs are overexpressed relative to their DNA copy number and this overexpression is significantly larger than that of non-TEP-TEs Wilcoxon rank-sum test, P=1.7e-07. **D)** Boxplot showing the ratios of X-linked versus autosomal polymorphic insertions for each TE in the TIDAL-fly (ref) database. TEP-TEs are depleted from the X compared to other TEs with polymorphic insertions. Wilcoxon rank-sum test, P = 0.029. **E)** Dot plot shows expression of selected piRNA pathway genes. Color of each dot corresponds to mean normalized and log-transformed expression within cell clusters. Dot size corresponds to the proportion of cells in each cluster expressing the marker.

Given that TEP-TEs are enriched on the Y chromosome, we next assessed whether their Y-linked copies are over-expressed in testes relative to their autosomal and X-linked copies. We used male and female WGS reads from *w1118* to identify male-specific (i.e. Y-linked) single-nucleotide variants in TEP-TEs. We then compared the relative abundance of each male-specific variant in testes RNA-seq data to its relative abundance in male WGS data (see Methods). A ratio larger than 1 indicates the presence of one or more Y-linked TE insertions that are expressed more highly than total-copy number alone would explain. For each expressed TE, we found the site of the male-specific allele most overexpressed relative to WGS depth. At these sites, male-specific TEP-TE alleles have significantly higher ratios of relative RNA to DNA coverage compared to male-specific alleles from non-TEP-TEs (Wilcoxon Rank-Sum test P=1.7e-07, Figure 5C). To confirm that this effect is specific to Y-linked TEP-TEs, we repeated our analysis using reference sites as well as autosomal (i.e. present in both males and females) variants. Contrary to the Y-linked TEP-TEs, these were expressed proportionately to WGS depth with no difference in expression proportion between TEP-TEs and other TEs (Wilcoxon Rank-Sum test P=0.24, Supplementary Figure 8F).

We next investigated whether the TEP-TEs show increased insertional activity in males. Polymorphic TE insertions reflect recent TE insertions that are still segregating within a population. If the TEP-TEs replicate more often in males compared to females, recent polymorphic insertions of these TEs should be depleted from the X chromosome because this chromosome is hemizygous in males and is therefore a smaller mutational target. We used the TIDAL-FLY database of polymorphic TE insertions (Rahman et al. 2015) for the Drosophila Genetic Reference Panel (DGRP) to compare insertion frequencies of TEP-TEs versus other active TEs. We found that non-TEP-TEs exhibit similar X and autosomal insertion rates across the DGRP lines whereas TEP-TEs exhibit a significantly reduced frequency of X-linked insertions relative to autosomal insertions, consistent with male-biased activity (Wilcoxon Rank-Sum test P= 0.029, Figure 5D).

*Ago3*, a piRNA pathway gene involved in the ping-pong piRNA amplification cycle, is present in germline stem cells and spermatogonia but undetectable in spermatocytes (Quénerch’du, Anand, and Toshie 2016). To determine whether there is a general trend of downregulation of piRNA pathway genes in spermatocytes compared to spermatogonia, we quantified expression of 31 piRNA pathway genes described in (Czech et al. 2018). We found a clear trend showing a striking downregulation of most piRNA pathway genes during the developmental transition from spermatogonia to spermatocytes (Figure 5E). Together, our results suggest that a burst of TE expression in Drosophila testes coincides with the activation of Y chromosome fertility genes and the downregulation of piRNA pathway genes.

## Discussion

In Drosophila ovaries, constrained developmental processes such as the nurse cell to oocyte mRNA transport pathway create a window of opportunity that TEs have evolved to exploit in order to increase their own copy numbers (Wang et al. 2018). Our results suggest a similar phenomenon has occurred in the testes, albeit via a different window of opportunity. A major source of TE activity in the testes is related to the presence of the Y chromosome itself. This chromosome acts as a safe harbor for TE insertions: The lack of recombination on the Y chromosome prevents efficient purging of Y-linked TEs from the population, allowing their accumulation along with other repetitive elements such as satellite DNA (Bachtrog 2013). However, the Y chromosome usually exists as tightly packaged, transcriptionally silent, heterochromatin. How can the presence of this inert chromosome lead to TE activation? Interestingly, there is evidence that the Y chromosome can act as a “sink” for heterochromatin: its presence may cause a genome-wide reallocation of repressive histone modifications, which can lead to TE de-repression (Henikoff 1996; Francisco and Lemos 2014; E. J. Brown and Bachtrog 2014; E. J. Brown, Nguyen, and Bachtrog 2020). On the other hand, Wei et al. have recently described a phenomenon that they term “Y toxicity” based on the upregulation of TEs present on the neo-Y chromosome of *Drosophila miranda* during embryogenesis (Wei, Gibilisco, and Bachtrog 2020). Their results suggest that transcription of the relatively large number of genes on the young neo-Y chromosome prevents complete silencing of this chromosome and therefore provides an opportunity for transcriptional activation of neo-Y-linked TEs.

Our results suggest that the Y toxicity phenomenon applies to older Y chromosomes as well. The ancient *Drosophila melanogaster* Y chromosome carries many fewer genes compared to the *D. miranda* neo-Y chromosome, however, at least six genes on the *D. melanogaster* Y chromosome are essential for male fertility (Brosseau 1960; Kennison 1981; Gatti and Pimpinelli 1983; Hazelrigg, Fornili, and Kaufman 1982). These genes are known as fertility factors and they are only expressed during spermatogenesis (Hardy, Tokuyasu, and Lindsley 1981). The three annotated fertility factors, *kl-2, kl-3*, and *kl-5*, each span as much as 4 Mb due to their extraordinarily large introns and become transcriptionally activated in primary spermatocytes, which coincides with a general decondensation and acetylation of the Y chromosome (Fingerhut, Moran, and Yamashita 2019). The transcription of three of these genes, *kl-5, kl-3*, and *ks-1*, is associated with the formation of large Y chromosome lampbrush loops (Bonaccorsi et al. 1988). The burst of TE expression that we describe here co-occurs with the activation of *kl-2*, *kl-3* and *kl-5*, as well as six other Y-linked genes: *ORY, ARY, Ppr-Y, Pp1-Y1, CG45765*, and *CCY*. Based on these results, we propose that the TEP-TEs have evolved to exploit a window of opportunity that occurs during the decondensation of the normally tightly packaged Y - linked chromatin, which is necessary for transcription of fertility factor genes. Notably, not all TE families with intact Y-linked insertions are members of the TEP, suggesting that additional features beyond Y-linkage, such as specific regulatory elements, are required for TEs to exploit this opportunity. Four TEP-TEs, *1731*, *412*, *copia*, and *micropia* have previously been shown to be highly expressed in primary spermatocytes in Drosophila and *micropia* transcripts are physically associated with Y chromosome lampbrush loops in *D. hydei* (Haoudi et al. 1997; Borie et al. 2002; Pasyukova et al. 1997; Lankenau, Corces, and Lankenau 1994).

Interestingly, another 20 TEP-TEs, including *gypsy*, have insertions located within the *flamenco* piRNA cluster. Chalvet et al identified multiple strains of *D. melanogaster* where active *gypsy* elements are confined to the Y chromosome (Chalvet et al. 1998). They proposed that Y-linked *gypsy* insertions are able to evade silencing by the ovary-dominant *flamenco* locus, which may explain the enrichment of *flamenco-regulated* TEs among members of the TEP. Indeed, more recent research has found that *flamenco-derived* piRNAs are almost an order of magnitude more abundant in ovaries compared to testes (P. Chen et al. 2020).

*Flamenco* is not unique in this respect – the majority of known piRNA clusters produce more abundant piRNA in ovaries compared to testes (P. Chen et al. 2020). Spermatocytes also lack a robust ping-pong amplification loop and the bulk of spermatocyte piRNAs come not from TEs, but rather two satellite repeats: *su(Ste)* and *AT-chX* (Nagao et al. 2010). Furthermore, piRNA factors such as Piwi and Ago3, while abundant in germline stem cells and spermatogonia, are missing or present at low levels in spermatocytes (Nagao et al. 2010; Quénerch’du, Anand, and Toshie 2016). Our analysis of scRNA-seq data confirm these findings (Figure 5E). Why do spermatocytes show a weakened piRNA response at a developmental timepoint when the TE-rich Y chromosome is de-repressed? One possibility is related to intragenomic conflict. Sex chromosomes are hotspots for genomic conflict (Bachtrog 2020) and small RNA pathways may play an outsize role in defending against meiotic drivers in the male germline (Courret et al. 2019; Lin et al. 2018). There is evidence that *stellate* and *su(Ste)* represent a cryptic meiotic drive system where X-linked *stellate* genes disrupt spermatogenesis and cause sex-ratio distortion in the absence of the Y-linked *su(Ste)* piRNA cluster (Hurst 1996; Bozzetti et al. 1995). The function of *AT-chX* is less clear. Although this locus was originally proposed to play a role in the developmental silencing of *vasa* during spermatogenesis (Nishida et al. 2007), recent results instead suggest of role for *AT-chX* in hybrid incompatibility (Kotov et al. 2019). Neither loci are present in the genomes of close relatives of *D. melanogaster*, suggesting that they are dispensable for spermatogenesis (Adashev et al. 2020; Kotov et al. 2019). The fact that both *su(Ste)* and *AT-chX* rapidly evolved to be essential for fertility in *D. melanogaster* is consistent with a role in mediating genetic conflict. This is especially clear for *su(Ste)* where the *stellate* protein is completely absent from wild-type flies (Adashev et al. 2020). If the *su(Ste)* and *AT-chX* piRNAs evolved to supress segregation distorters or other forms of selfish elements, it would suggest that there has been a tradeoff in the piRNA system in *D. melanogaster* spermatocytes, where increased abundance of *su(Ste)* and *AT-chX* piRNAs comes at a cost of impaired TE silencing. Future work investigating the peculiarities of TE silencing in the testes will help shed light upon this and other constraints imposed by the various roles of piRNAs in the male germline, including host gene regulation, TE silencing, and the resolution of intragenomic conflicts.

## Methods

All code is provided as a snakemake workflow (Mölder et al. 2021) at github.com/Ellison-Lab/TestisTEs2021. Male and female *w1118* whole genome sequencing data and *w1118* total RNA-seq data are deposited at PRJNA727858.

### Repeat masking and custom reference sequence generation

All repeat masking was performed with RepeatMasker (Smith, Hubley, and Green 2013) with the following options: “-e ncbi -s -no_is -nolow.” We used RepBase *D. melanogaster* consensus TE sequences (version 20170127) (Bao, Kojima, and Kohany 2015) as a custom library.

For the purposes of generating reference sequences for alignments, we appended the consensus TE sequences to the masked *D. melanogaster* r6.22 sequence.

### scRNA-seq processing

Single cell RNA-seq data was downloaded from PRJNA548742 and PRJNA518743. We used 10X Genomics cellranger software to align and quantify the data (Zheng et al. 2017). We generated a cellranger index from the previously described custom reference sequence using cellranger’s “mkref” command with default parameters. We aligned scRNA-seq reads using cellranger’s “count” command with default parameters. We used cellranger’s filtered count matrices for further analysis.

We first summed counts assigned to the LTR and internal sequences of class I LTR retrotransposons. For each scRNA-seq replicate, we next applied scrublet v0.2.1 (Wolock, Lopez, and Klein 2019) to these unnormalized count matrices to identify and filter putative heterotypic doublets. We used scanpy v1.6.0 (Wolf, Angerer, and Theis 2018) to retain genes detected in at least 3 cells and then cells with at least 250 and fewer than 5000 detected genes. We removed cells with more than 5% of remaining UMIs assigned to mitochondrion-encoded genes. We normalized UMI counts to 10000 per cell and applied log transformation with a pseudo-count of 1. We identified highly variable genes using scanpy’s “highly_variable_genes” method with default parameters. We next scaled counts using scanpy’s “scale” method and applied scanpy’s “regress_out” method to remove count variance associated with cell cycle and mitochondrial UMI counts.

For each replicate, we used scanpy to perform principal component analysis on highly variable host genes and calculate nearest neighbor graphs using 15 principal components and 25 neighbors. We called cell clusters using the Leiden algorithm (Traag, Waltman, and van Eck 2019) via scanpy with a resolution parameter of 0.35. We combined all three larval scRNA-seq replicates using scanpy’s “ingest” method.

Automated cell type assignment was performed using Garnett v0.2.17 (Pliner, Shendure, and Trapnell 2019) and a set of curated marker genes (Supplementary Table 1).

### Consensus ICA for GEP Detection

We chose ICA to identify gene expression programs because it performs highly with respect to recovering known functional gene modules and because it is easily adaptable to finding partially overlapping modules (Saelens, Cannoodt, and Saeys 2018). We standardized the normalized, log-transformed expression matrix to have zero mean and unit variance. Standardized scores were clipped to a maximum absolute value of 10.

To generate stable modules that are robust to stochastically varying ICA solutions, we applied a consensus approach previously applied to non-negative matrix factorization gene module detection (Kotliar et al. 2019). We used FastICA via sklearn (Pedregosa et al. 2012) to decompose the standardized expression matrix into 90 components 100 times, then concatenated the resulting gene *x* module matrices, partitioned all modules into 90 clusters using k-means clustering, and averaged the per-cell scores within each partition to yield a consensus cell *x* module matrix. Within the same partitions, we averaged per-gene scores from cell *x* module matrices to generate a consensus cell *x* module matrix.

We assigned genes to each program by applying fdrtool (Strimmer 2008) to the vector of gene weights for each module. Genes with FDR q-values less than 0.005 for each module were considered members of the module.

### GEP parameter optimization

Use of ICA or other matrix decomposition approaches for gene program detection requires *a priori* assumptions about the optimal number of components (*k*) to request from the decomposition algorithm. Additionally, generation of discrete gene lists for each gene program requires application of arbitrary score cutoffs to determine program membership for each gene.

To reduce bias and use of arbitrary cutoffs, we used a grid search approach to choose *k* and the q-value cutoff for membership. Briefly, we ran consensus ICA in triplicate for combinations of q-value cutoffs between 0.005 and 0.1 and *k* between 20 and 120. We performed pathway enrichment analysis for each program discovered in each consensus ICA replicate and for each run calculated the percentage of GO:BP terms with significant enrichment as well as the percentage of programs in each run that show a unique significant enrichment. We then rescaled these scores to a maximum of 1 and calculated a joint score by multiplying them together. For our final set of gene programs, we ran consensus ICA a final time with the *k* and q-value that maximized the average joint score across all three test replicates.

### Poly-A RNA-seq

We trimmed poly-A selected RNA-seq (SRR7276830, SRR7276831, SRR7276832, SRR7276833) with fastp v0.20.0 (S. Chen et al. 2018) and aligned to the custom reference using STAR v2.7.3 (Dobin et al. 2013) with chimeric junction detection turned on and “--chimScoreJunctionNonGTAG 0”. Other non-default parameters used are available via the linked github repository.

We calculated normalized coverage for each strand using deeptools v3.3.1 (Ramírez et al. 2014) “bamCoverage” command with “--smoothLength 150.”

### WGS library preparation

20 0- to 3-day old *w1118* males or females were collected on dry ice and then homogenized using an electric pestle. Qia-Amp DNA Micro kit was used according to instructions. DNA was diluted to 40 ng/ul in 55 ul of Elution Buffer and sheared in a Covaris sonicator with settings as follows: 10% duty cycle, 2.0 intensity, 200 cycles per burst, 1 cycle, 45 second process time.

WGS library generation protocol was adapted from the Marshall Lab DamID-seq protocol available at marshall-lab.org (Marshall et al. 2016). Briefly, sheared DNA was purified with homemade purification beads. End repair was performed with T4 DNA Ligase (NEB M0202S), T4 DNA Polymerase (NEB M0203S), PolI Klenow fragment (NEB M0210S), T4 Polynucleotide kinase (NEB M0201S). Adenylation was performed with 3’-5’ Klenow Fragment (NEB M0212L). Adaptors were ligated with NEB Quick Ligase for 10 minutes at 30°C before two rounds of cleanup with homemade beads. NEBNext UltraII Q5 kit (NEB M0544) was used for PCR enrichment. A final round of cleanup with homemade beads was performed before quantification and sequencing.

### WGS processing

We trimmed reads using cutadapt v3.2.0 (Martin 2011) with options “-q 20 -m 35.” We aligned trimmed reads with bwa-mem2 v2.0 (Vasimuddin et al. 2019), removed duplicate reads with picard v2.22.1 (“Picard” n.d.) with option “VALIDATION_STRINGENCY=LENIENT”, and filtered out multimappers with samtools v1.10 (Danecek et al. 2021).

To estimate TE copy number estimation we used mosdepth v0.3.1 (Pedersen and Quinlan 2018) to calculate genome-wide read coverage in 100 bp bins, then compared TE coverage to autosome coverage.

We identified male-specific polymorphic sites with Rsamtools (Morgan M, Pagès H, Obenchain V, Hayden N 2020) by finding mismatches with a base quality of at least 10 and at least 15 supporting male reads but lacking supporting female reads.

### Total RNA-seq library preparation

We used approximately 100 pairs of testes from 3-5-day old mated *w1118* males. The testes were dissected in 1X PBS and transferred into 200 μL RNAlater Solution. Tissue was pelleted by centrifuging at 5000g for 1 min at 4 °C. Supernatant was removed and 300 μL 1x DNA/RNA Shield was added before homogenization with an electric pestle. Homogenized tissue was digested with Proteinase K at 55 °C for at least 30 min. RNA was purified with the Zymo Quick-RNA Plus Kit (R1057).

Using up to 5 μg total RNA, ribosomal RNAs were removed suing iTools rRNA depletion Kit from Galen Laboratory Supplies (dp-P020-000007) and Thermo Fisher MyOne Streptavidin C1 Dynabeads (#65001). RNA Clean & Concentrator-5 kit from Zymo Research (R1015) was used to purify rRNA-depleted RNA. Starting with 1 ng-100 ng purified rRNA-depleted RNA, Illumina libraries were generated using NEBNext Ultra II Directional RNA Library Prep Kit for Illumina (E7760).

### Total RNA-seq processing

Raw reads were trmmed with fastp v0.20.0. We used STAR v2.7.5 (Dobin et al. 2013) to align total RNAseq reads to a bait reference composed of Flybase release 6.22 tRNA sequences and miscRNA sequences. We then aligned unmapped reads to our custom reference and provided STAR with a VCF file containing male-specific variants.

### DGRP Polymorphic TE insertions

Using the TIDAL-Fly polymorphic TE insertion database (Rahman et al. 2015), we found the number of unique polymorphic insertions on the X chromosome and on autosomes, excluding chromosome 4, across the Drosophila Genome Reference Panel for all TEs in our custom reference. For all TEs with at least 1 X-linked and 1 autosomal insertion among all DGRP lines, we calculated the ratio of X-linked insertions per megabase to autosomal insertions per megabase.

### RNA-FISH

Custom Stellaris FISH probes recognizing *EAChm* labeled with Quasar670 and against Accord2 and Quasimodo2 labeled with CAL Fluor Red 610 were designed using Stellaris’ probe design tool available at www.biosearchtech.com/stellarisdesigner. Default parameters were used for *EAChm* probes. Probes against Accord2 and Quasimodo2 were designed with masking parameter 2. To ensure specificity of the resulting probes, we used BLAST (Camacho et al. 2009) to align to consensus TE sequences used for masking and custom reference generation to ensure that all probes show complementarity to their intended target only. *Accord2* and *Quasimodo2* probes were also blasted against *Drosophila melanogaster* REFSEQ sequences and any individual probes with more than 16 nucleotide matches to another sequence were removed from the final probe set.

Strain *w1118* flies maintained at room temperature were mated for 4 hours and offspring were grown at 25°C until reaching the third instar. We dissected L3 males in sterile 1X PBS and fixed testis in 3.7% formaldehyde solution at room temperature for 45 minutes. Testis were washed twice with 1X PBS and submerged in 70% ethanol at 4°C overnight. Hybridizations were carried out according to instructions available on the manufacturer website.

Image slices were captured on a Carl Zeiss LSM880 AxioObserver with a C-Apochromat 40x/1.2 W Korr FCS M27 water immersion objective. 2D deconvolution was performed using ZEN Black software. Further contrast adjustments and image overlays were performed with Fiji (Schindelin et al. 2012).

## Acknowledgements

The authors acknowledge the laboratory of Dr. Maureen Barr for use of their confocal microscope and Dr. Juan Wang for microscopy training and assistance. The authors also acknowledge the Office of Advanced Research Computing (OARC) at Rutgers, The State University of New Jersey for providing access to the Amarel cluster and associated research computing resources that have contributed to the results reported here.

---

**Supplementary Table 1.**

|  |  |  |
| --- | --- | --- |
| aly | Spermatocyte | White-Cooper et al. 1998; Mahadevaraju et al. 2021 |
| sa |  | White-Cooper et al. 1998; Mahadevaraju et al. 2021 |
| tbrd-1 |  | Theofel et al. 2014; Leser et al. 2012; Mahadevaraju et al. 2021 |
| tbrd-2 |  | Theofel et al. 2014; Mahadevaraju et al. 2021 |
| bol |  | Maines and Wasserman 1999; Mahadevaraju et al. 2021 |
| fzo |  | Hwa et al. 2002; Mahadevaraju et al. 2021 |
| Ance |  | Hurst et al. 2003; Mahadevaraju et al. 2021 |
| dj |  | Santel et al. 1997; Lim, Tahrayrah, and Chen 2012; Mahadevaraju et al. 2021 |
| ocn |  | Mikhaylova and Nurminsky 2011; Mahadevaraju et al. 2021 |
| CycB |  | Perezgasga et al. 2004; Mahadevaraju et al. 2021 |
| twin |  | Mahadevaraju et al. 2021 |
| bb8 |  | Vedelek et al. 2016; Mahadevaraju et al. 2021 |
| AGO3 | Spermatogonia | Quenerch' du et al. 2016; Mahadevaraju et al. 2021 |
| vas |  | Lasko and Ashburner 1990; Mahadevaraju et al. 2021 |
| bam |  | Eun et al. 2013; Mahadevaraju et al. 2021 |
| aub |  | Quenerch' du et al. 2016; Mahadevaraju et al. 2021 |
| p53 |  | Monk, Abud, and Hime 2012; Mahadevaraju et al. 2021 |
| Dek |  | Mahadevaraju et al. 2021 |
| osa |  | Mahadevaraju et al. 2021 |
| CycA |  | Mahadevaraju et al. 2021 |
| tj | Cyst | Li et al. 2003; Mahadevaraju et al. 2021 |
| eya |  | Zoller and Schulz 2012; Mahadevaraju et al. 2021 |
| piwi |  | Nishida et al. 2007; Mahadevaraju et al. 2021 |
| bnb |  | Terry et al. 2006; Mahadevaraju et al. 2021 |
| Nrt |  | Terry et al. 2006; Mahadevaraju et al. 2021 |
| Wnt4 |  | Terry et al. 2006; Mahadevaraju et al. 2021 |
| Fas3 | Terminal Epithelial | Mahadevaraju et al. 2021; Mahadevaraju et al. 2021 |
| nord |  | Mahadevaraju et al. 2021; Mahadevaraju et al. 2021 |
| Piezo |  | Mahadevaraju et al. 2021; Mahadevaraju et al. 2021 |
| Sox100B | Pigment | Nanda et al. 2009; Mahadevaraju et al. 2021 |
| ems |  | Nanda et al. 2009; Mahadevaraju et al. 2021 |

**Supplementary Figure 1.**
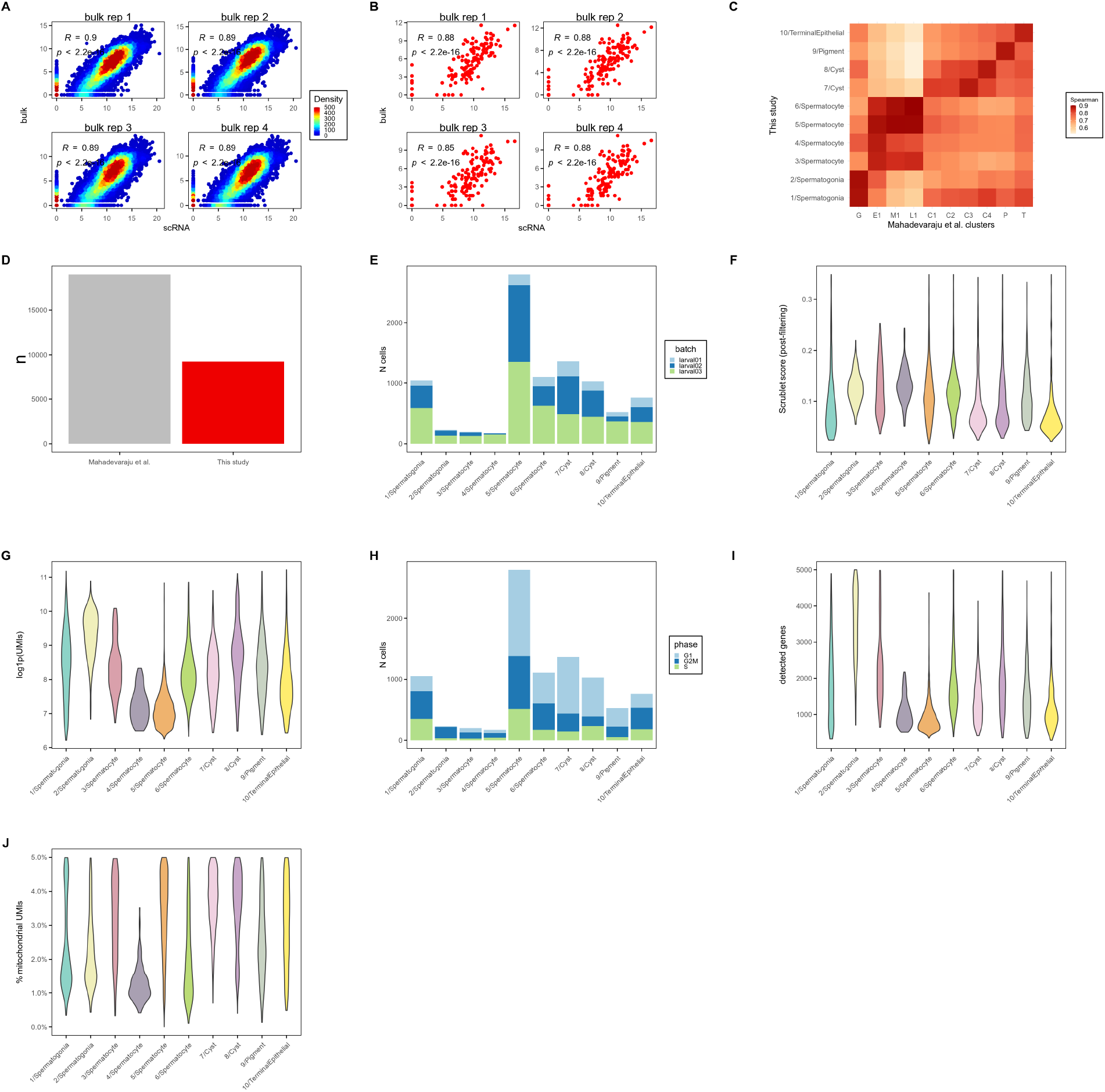
scRNA-seq data. **A)** Scatterplots show pseudo-bulk expression derived from our *w1118* scRNA pipeline (see **Methods**) versus bulk expression for 4 *w1118* testis poly-A RNA-seq replicates generated by Mahadevaraju et al. 2021 (see **Methods**). Each replicate shows strong correlation with pseudo-bulk (all Pearson’s R >= 0.89, P<2.2e-16). **B)** Scatterplots show same analysis described in **A** but restricted to TEs (all Pearson’s R >= 0.85, P<2.2e-16). **C)** Heatmap shows correlation between scRNA-seq expression estimates derived from our pipeline for all clusters identified in this study compared to clusters identified by Mahadevaraju et al. 2021 (source of data). **D)** Barplot shows the number of cells used in Mahadevaraju et al. 2021 (gray) and this study (red). **E)** Barplot shows the number of cells assigned to each cluster. Bars are divided and colored by the scRNA replicate from which cells are derived. **F)** Distribution of doublet scores (see **Methods**) for cells in each cluster after all filtering steps. **G)** Distribution of total log1p(UMIs) for cells in each cluster after all filtering steps. **H)** Barplot shows the number of cells assigned to each cluster. Bars are divided and colored by predicted cell cycle phase (see **Methods**). **I)** Distribution of total detected genes for cells in each cluster after all filtering steps. **J)** Distribution of the percentage of total pre-filtering mitochondrial reads for cells in each cluster after all filtering steps.

**Supplementary Figure 2.**
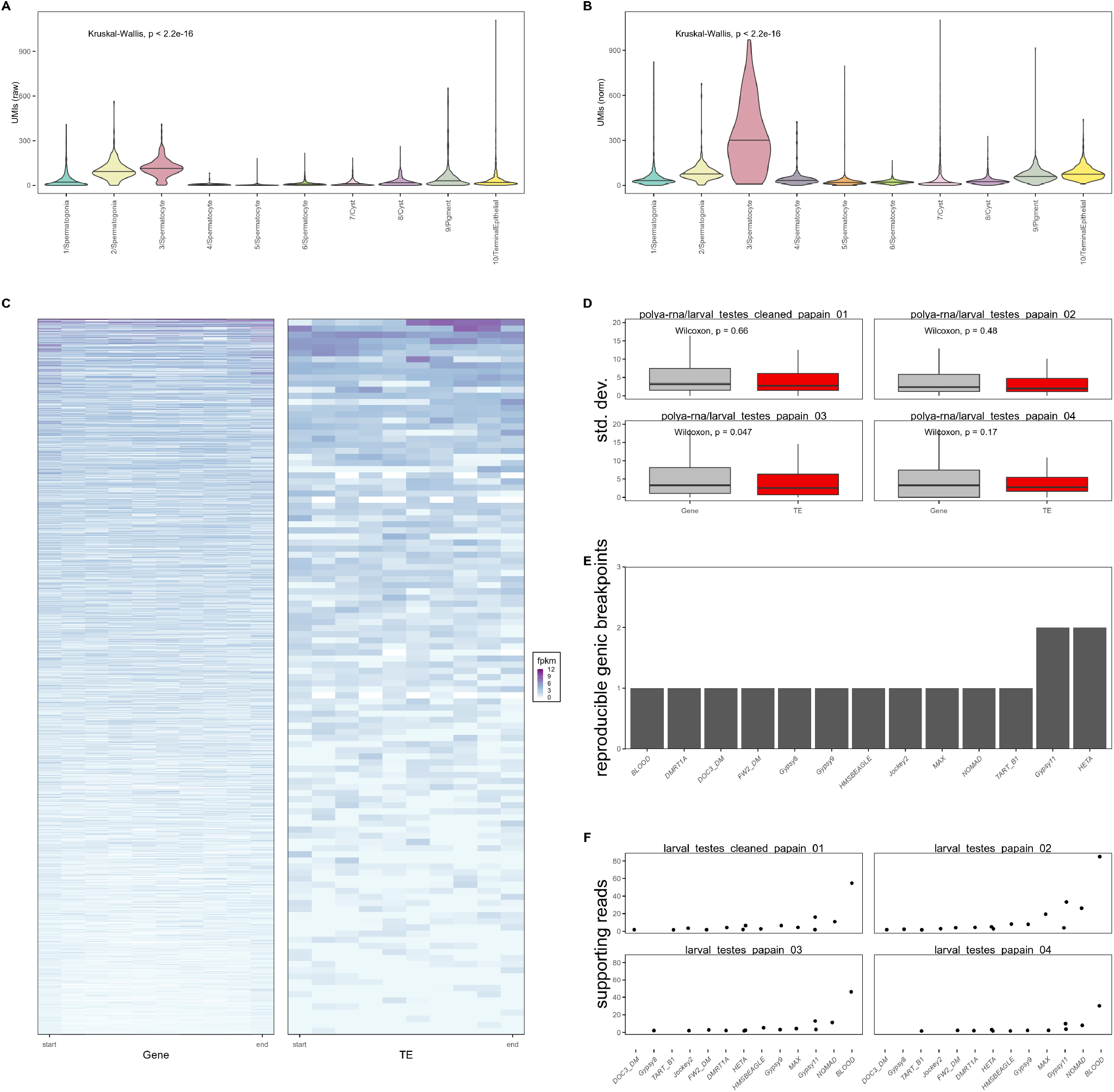
TE expression in scRNA. **A**) Violin plots show distributions of raw TE-mapping UMI counts in each L3 *w1118* cell cluster. TE counts vary significantly among the clusters (Kruskal-Wallis, p < 2.2e-16). **B**) Violin plots show distributions of depth normalized TE-mapping UMI counts in each L3 *w1118* cell cluster. TE counts vary significantly among the clusters (Kruskal-Wallis, p < 2.2e-16). **C)** Heatmaps show distribution of sense-strand poly-A RNA-seq signal for single-isoform host gene mRNAs (left) and detected TEs (right) in bins comprising the full length of the respective features. poly-A RNA-seq data generated by Mahadevaraju et al 2021 (see **Methods**). **D)** Boxplots show standard deviations of expression across bins for host genes (gray) and TEs (red). Three of four replicates show no significant difference in variability of poly-A signal across bins within features (Wilcoxon rank-sum test P>0.05). Replicate 3 shows a significant difference (Wilcoxon rank-sum test, P=0.047). **E)** Bar plot shows number of genic fusions reproducibly found in poly-A RNA-seq data. **F)** For each TE introduced in **E**, the y-axis position of each point represents the number of uniquely-mapping chimeric reads detected by STAR that support each breakpoint (see **Methods**).

**Supplementary Figure 3.**
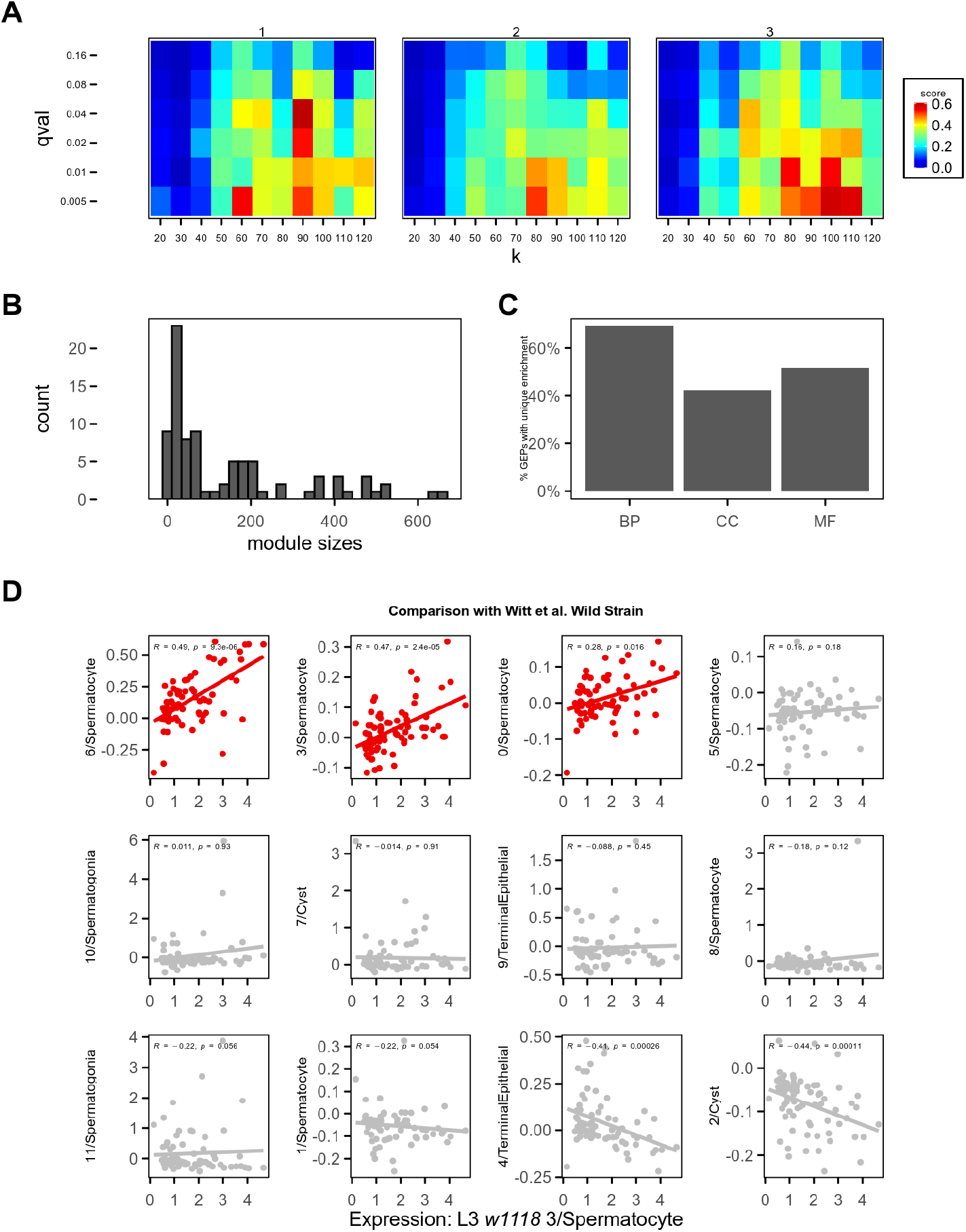
Gene Expression Program detection via consensus approach to Independent Component Analysis. **A)** Heatmaps correspond to independent replicates of the grid search approach used to optimize ICA component number (*k*) and *q*-value cutoff. Color intensity corresponds to the enrichment score (see **Methods**) for each combination of *q* and *k*. **B)** Histogram shows module sizes among the set of GEPs used in main analysis. The majority of modules detected include fewer than 100 features. **C)** Barplots show percentage of discovered GEPs with at least 1 unique Biological Process, Cellular Component, or Molecular Function enrichment. **D)** Scatterplots show relationship of TE expression in *w1118* Cluster 3/Spermatocyte versus all clusters detected by our pipeline in the “Wild Strain” dataset generated by Witt et al. 2019. Multiple spermatocyte clusters (highlighted in red) show TE expression patterns similar to 3/Spermatocyte.

**Supplementary Figure 4.**
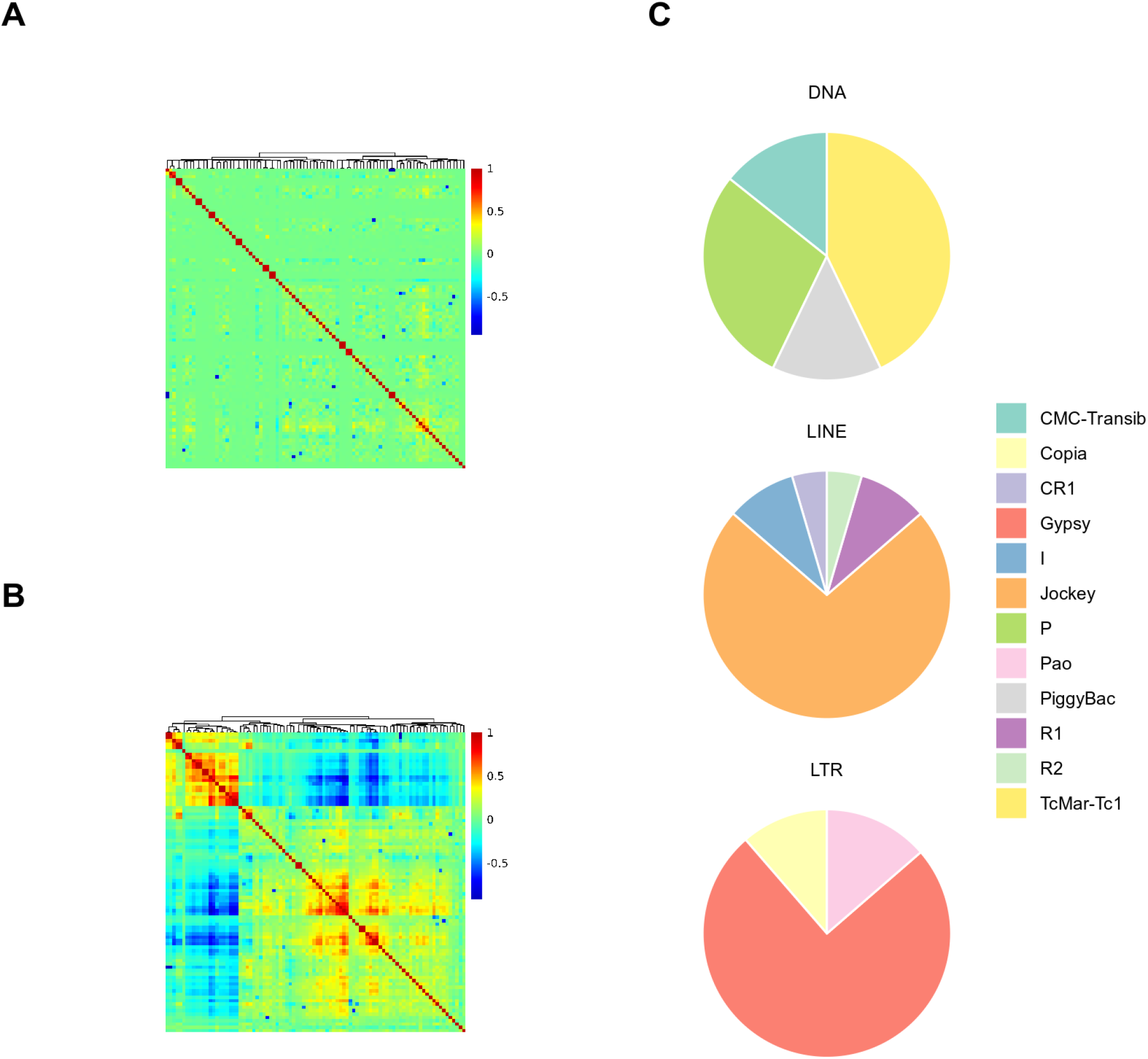
Redundancy and TE content of GEPs. **A)** Clustering of GEPs by cell usage score (consensus ICA source matrix). **B)** Clustering of GEPs by gene membership score (consensus ICA mixing matrix). For **A,B** `1-Pearson’s R` is used as a distance metric. **C)** Breakdown of specific DNA TE, LINE, and LTR TE families in GEP-27.

**Supplementary Figure 5.**
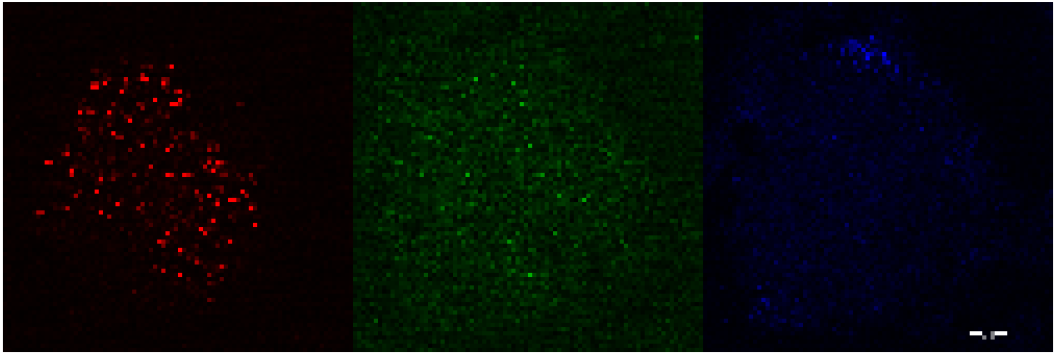
ACCORD2 *and* EAChm *expression in L3 testes*. Representative slice of multiplexed RNA-FISH in whole-mount 3rd larval instar w1118 testis. Image is split by color channel. *ACCORD2* and *EAChm* expression is detected in the middle region of the testis, where primary spermatocytes are located. Red: *EAChm*; green: *ACCORD2*; blue: DAPI.

**Supplementary Figure 6.**
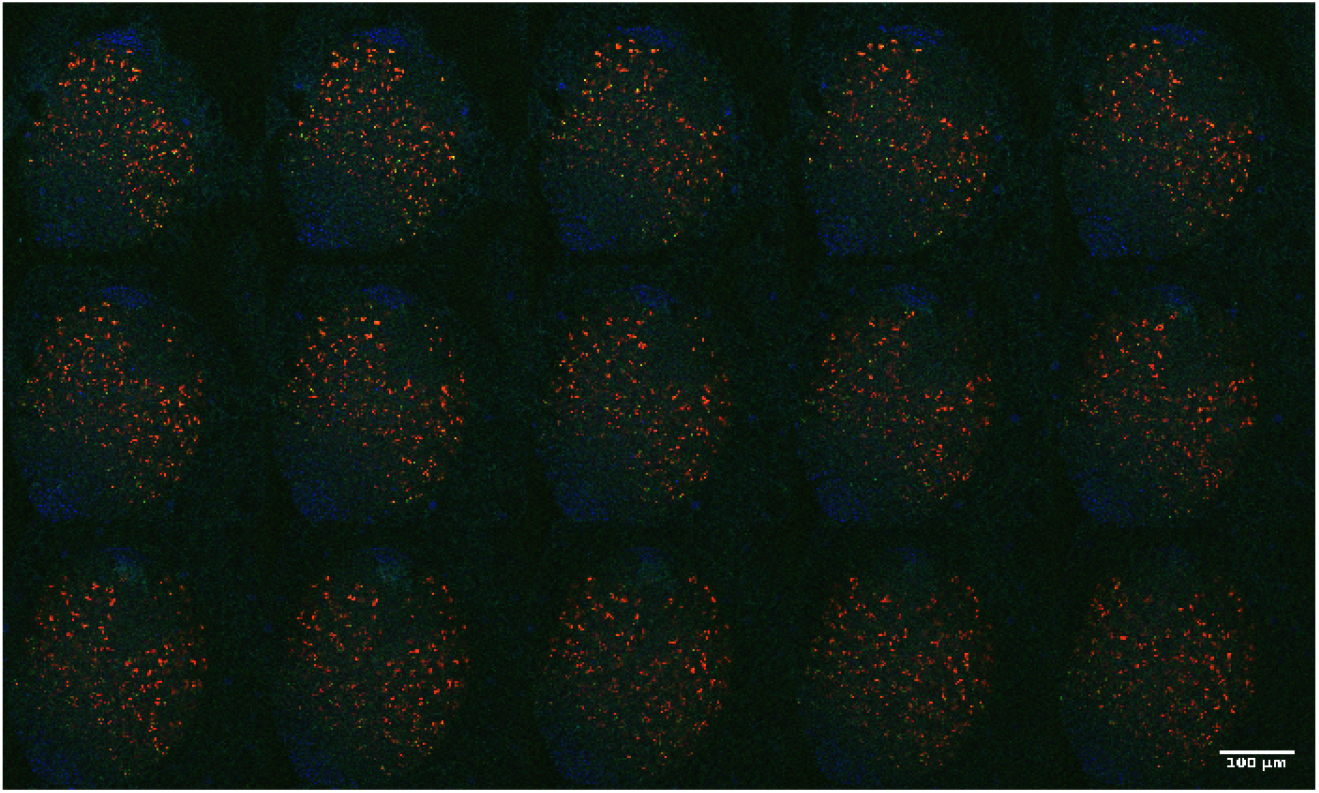
ACCORD2 *and* EAChm *expression in L3 testes*. Z-slices montage of multiplexed RNA-FISH in whole-mount 3rd larval instar w1118 testis. *ACCORD2* and *EAChm* expression is detected in the middle region of the testis, where primary spermatocytes are located. Red: *EAChm*; green: *ACCORD2*; blue: DAPI.

**Supplementary Figure 7.**
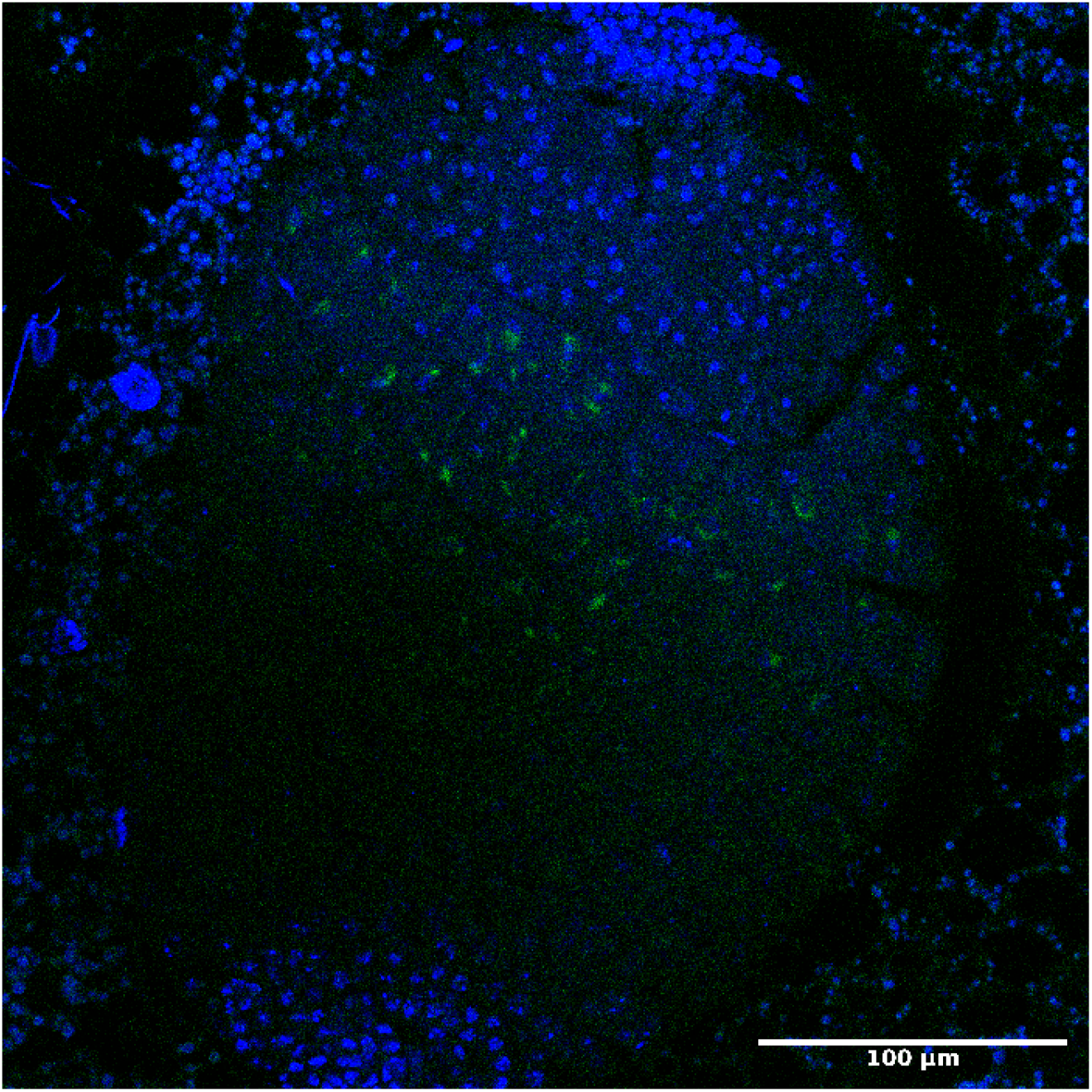
QUASIMODO2 *expression in L3 testes*. Representative RNA-FISH z-slice in whole-mount 3rd larval instar *w1118* testis. *QUASIMODO2* expression is detected in the middle region of the testis, where primary spermatocytes are located. Green: *QUASIMODO2*; blue: DAPI.

**Supplementary Figure 8.**
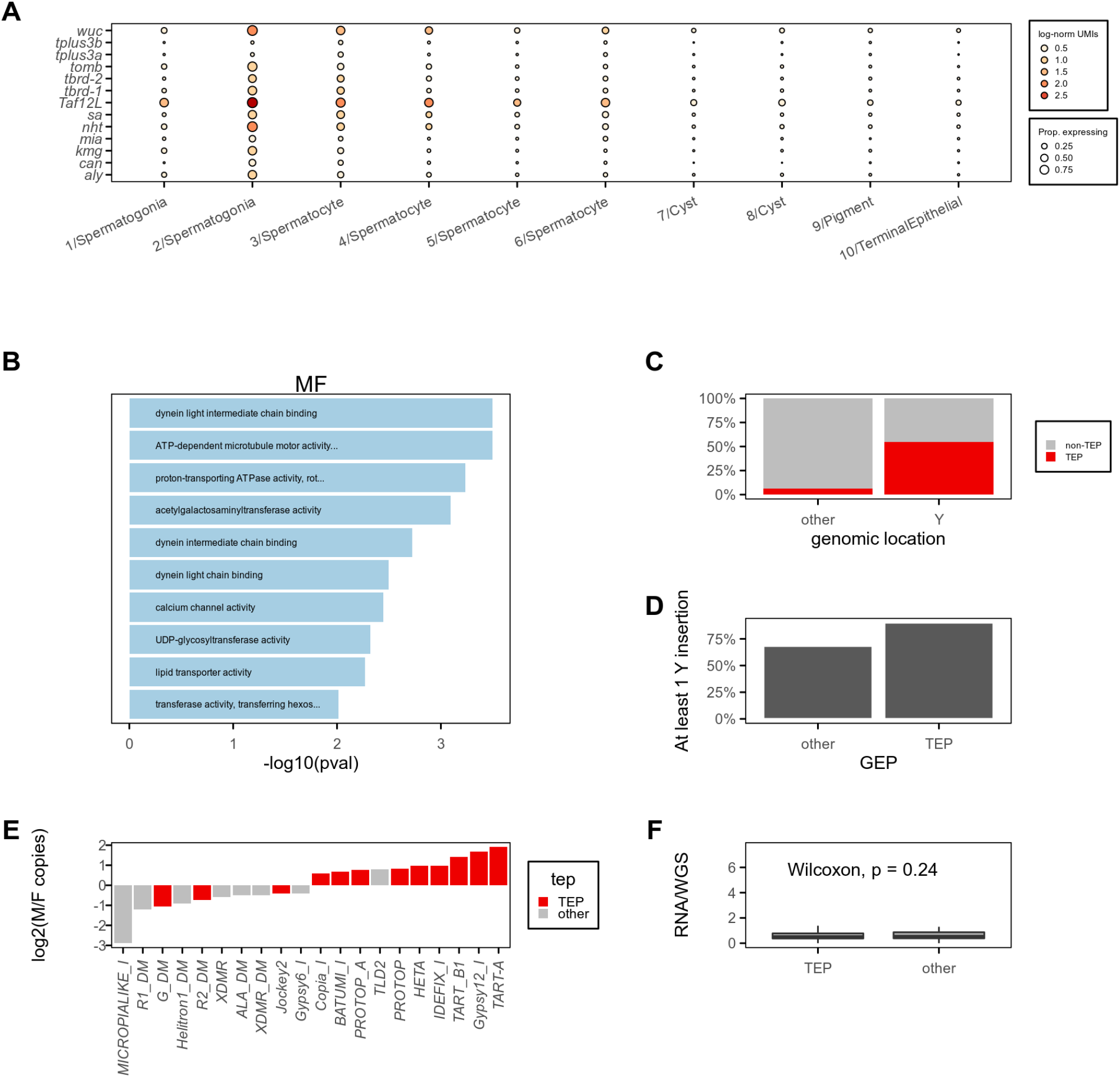
TEP-TEs are enriched on the Y-chromosome. **A)** Dot plot shows expression of selected effectors of spermatocyte transcriptional programs. Color of each dot corresponds to mean normalized and log-transformed expression within cell clusters. Dot size corresponds to the proportion of cells in each cluster expressing the marker. **B)** Bars show strength of top 10 enriched Gene Ontology Molecular Function terms for GEP-27. **C)** Bar plot shows proportion of Y chromosome genes or other genes that are assigned to the TE-enriched Program or other GEPs. Chi-square test P=1.7e-05. **D)** Barplot shows percentage ofTEP or non-TEP TEs with at least 1 Y-linked insertion detected by RepeatMasker in the heterochromatin-enriched assembly described by Chang and Larracuente 2019. **E)** Bars represent the ratios of male to female estimated copies for the top 10 male- and female-enriched TEs. Bars are colored by membership in TEP. **F)** Allele specific analysis of TE expression (see **Methods**) shows that non-Y-linked copies of TEP TEs are expressed proportionately to their DNA copy number. Wilcoxon rank-sum test, P=0.24.

